# Aerobic methane production by methylotrophic *Methylotenera* in groundwater

**DOI:** 10.1101/2024.07.22.604536

**Authors:** Shengjie Li, Xiaoli Dong, Pauline Humez, Joanna Borecki, Jean Birks, Cynthia McClain, Bernhard Mayer, Marc Strous, Muhe Diao

## Abstract

*Methylotenera* are signature denitrifiers and methylotrophs commonly found together with methanotrophic bacteria in lakes and freshwater sediments. Here we show that three distinct *Methylotenera* ecotypes were abundant in methane-rich, Pleistocene-aged groundwaters. Just like in surface water biomes, groundwater *Methylotenera* often co-occurred with methane-oxidizing bacteria, even though they were generally unable to denitrify. One abundant *Methylotenera* ecotype expressed a pathway for aerobic methane production from methylphosphonate. This phosphate-acquisition strategy was recently found to contribute to methane production in the oligotrophic, oxic upper ocean. Gene organization, phylogeny and 3D protein structure of the key enzyme, C-P lyase subunit PhnJ were consistent with a role in phosphate uptake. We conclude that phosphate may be a limiting nutrient in productive, methane rich aquifers and that methylphosphonate degradation can contribute to groundwater methane production.

## Introduction

*Methylotenera* (*Pseudomonadota*, *Gammaproteobacteria*, *Burkholderiales*, *Methylophilaceae*) are commonly found in freshwater lakes^1–3^. They are methylotrophs and facultative denitrifiers, specialized in using reduced one-carbon compounds such as methanol and methylamine as their carbon and energy sources^4^. Several isolates were obtained from Lake Washington sediments, including *Methylotenera mobilis* JLW8^5^, *Methylotenera* Strain G11 and L2L1^1^, *Methylotenera versatilis* 301 and 30S^6^. *Methylotenera oryzisoli* La3113 was isolated from rice field soil^7,8^.

Methane is a potent greenhouse gas that is also commonly found in groundwater^9–11^. Although *Methylotenera* do not directly use methane, they are known to engage in partnerships with methane-oxidizing *Methylobacter*. In these partnerships, which are beneficial to both, *Methylobacter* feed methanol and formaldehyde to denitrifying *Methylotenera*^12–16^.

*Methylotenera* have mainly been studied in surface water, but a recent study found they were one of the most abundant groundwater residents (up to 88% of all bacteria) ^17,18^. Fifty-five amplicon sequence variants (ASVs) affiliated with *Methylotenera* were found in groundwater from 138 wells, indicating that these bacteria are even more common and diverse in groundwater than in surface water. These two habitats differ in important ways. Compared to surface water, opportunities for migration of microorganisms in groundwater are more limited. Groundwater can be over 10,000 years old, predating the last deglaciation and the start of the Holocene, and many aquifers are “confined”, covered by an impermeable layer that prevents mixing with younger water^19^.

Groundwater aquifers are also structured habitats. The occurrence of spatial gradients might be needed to explain apparently conflicting observations. For example, a single groundwater sample may contain both oxygen and abundant aerobic *Methylotenera* as well as methane and anaerobic, methane-producing *Methanobacteriaceae*^17^. A similar paradox was observed in the ocean, where aerobic methane production from organic phosphonates finally explained the presence of methane in an oxic ecosystem^20–22^. Here we show that organic phosphonates may also contribute to methane production in pristine groundwater aquifers.

Briefly, we recovered thirty-six metagenome-assembled-genomes (MAGs) of *Methylotenera* from nineteen groundwater samples, with methane concentrations varying from below detection limit to 69 mg/L. We investigated the ecophysiology and biogeography of these *Methylotenera* populations based on their genomes and protein expression. Three groundwater ecotypes of *Methylotenera* were observed. One of the ecotypes appeared to produce methane aerobically from methylphosphonate.

## Materials and Methods

### Groundwater sampling, amplicon, metagenome and proteome sequencing

Sampling of groundwater wells in Alberta, Canada, and measurements of physico-chemical parameters have been previously described^18^. Detailed information of well locations, aquifers, lithology, geology and aqueous geochemistry of the groundwater samples is provided in Supplementary Table S1. DNA extraction, amplicon sequencing, shotgun metagenomic sequencing, protein extraction and analysis were previously described^17^. This study included an amplicon sequencing dataset with 265 groundwater samples collected from 138 wells from 2016 to 2022, 26 metagenomes collected from 25 wells in 2019 and 2022, and 5 proteomes collected from 5 wells in 2019.

### Metagenome and proteome analysis

Quality trimming of raw reads, assembly, binning and read mapping were previously described^17^. Quality and characteristics of the obtained MAGs were estimated with CheckM2 v0.1.3^23^. Estimated genome size was calculated as actual MAG size divided by estimated MAG completeness. The taxonomic identity of MAGs was obtained with GTDBtk v2.3.0^24^. In total, thirty-six MAGs affiliated with *Methylotenera* were obtained. There were twenty high-quality MAGs (>90% completeness and <5% contamination), thirteen medium-quality MAGs (>50% completeness and <10% contamination) and three MAGs of lower quality (completeness >85% and 10%-15% contamination). Average nucleotide identity between the thirty-six MAGs was obtained with FastANI v1.33^25^. The twenty high-quality MAGs were used to infer the metabolism based on gene content. Expression of proteins of nine *Methylotenera* MAGs was detected in proteomes obtained for five proteomes as previously described^17^.

### Relative abundance of populations

The relative abundance of *Methylotenera* populations associated with MAGs in each metagenome was estimated with the “checkm coverage” and “checkm profile” commands within CheckM v1.1.3b^26^. The relative abundance of methanogens was calculated in two ways: (1) based on sequencing depths of MAGs, and (2) based on sequencing depths of 16S rRNA genes reconstructed by phyloFlash^27^. The abundances calculated with the two approaches were in good agreement (Supplementary Table S1).

### Annotation

MAGs and unbinned contigs were annotated with MetaErg v2.3.39^28^. Key enzymes involved in methylotrophic processes were analyzed with Hidden Markov Models (HMMs) obtained from a previous study^29^, which collected 1016 reference protein sequences from published genomes of methylotrophs. Twelve enzymes involved in cobalamin synthesis and transport were analyzed with HMMs downloaded from TIGRFAM and Pfam as previously described^30^.

### Estimation of the amount of time passed since the common ancestor of closely related *Methylotenera* colonized these groundwaters

Among *Methylotenera* MAGs, there were several that were very closely related, with average nucleotide identities of 99.906% to 99.999%. To estimate the amount of time that may separate these close relatives found in thousands-year-old groundwater, we made use of previous estimates of bacterial mutation rates. The bacterial mutation rate varies from 10^-5^ to 10^-8^ nucleotide substitutions per site per year^31,32^. The mutation rate decreases as the sampling timescale increases, because a substantial fraction of mutations may appear over a short time scale but will ultimately be removed by natural selection. Therefore, the lower rate of 10^-8^ nucleotide substitutions per site per year was used.

### Phylogeny of the *Methylotenera* genus

Eighty-eight *Methylotenera* reference genomes together with 313 genomes from other genera of the *Methylophilaceae* family were downloaded from the Genome Taxonomy Database (GTDB) version 214^24^. Thirty-six *Methylotenera* MAGs recovered from this study together with these reference genomes were included in the phylogenetic analysis. Four genomes from other families in the *Burkholderiales* order were used as the outgroup. 71 single-copy marker genes included in the bacterial marker gene set in Anvi’o v7.1^33^ were extracted from the genomes and aligned with the “anvi-get-sequences-for-hmm-hits” command. The phylogenetic tree was constructed with the “anvi-gen-phylogenomic-tree” command which used FastTree^34^ and visualized using iTOL v6.9 (https://itol.embl.de/).

### Phylogeny, gene organization and 3D protein structure analysis of the C-P lyase core complex subunit J

A maximum-likelihood phylogenetic tree of the C-P lyase core complex subunit J (PhnJ) was constructed. First, PhnJ sequences from both binned and unbinned contigs in groundwater metagenomes were extracted based on annotation with MetaErg v2.3.39^28^ and reference sequences were obtained from the Metaerg database. This database contains RefSeq-annotated proteins from organisms in the GTDB database version 214^24^. Reference PhnJ sequences were dereplicated and proteins shorter than 150 amino acids were not further considered. Next, a multiple sequence alignment of the recovered PhnJ sequences and reference sequences was constructed using MAFFT v7.508 with default parameters^35^. Poorly aligned regions were removed with trimAl v1.4^36^. The phylogenetic tree was constructed with “-m MFP-B 1000” in IQ-TREE v2.2.0.3^37^ and visualized using iTOL v6.9 (https://itol.embl.de/).

The gene clusters of the Phn complex found in MAGs were compared with the experimentally verified *Escherichia coli* K-12 gene cluster^38^ with clinker v0.0.28^39^.

3D protein structure of the PhnJ proteins in *Methylotenera* and *Methylomonadaceae* was predicted with ColabFold v1.5.5^40^ by combining the fast homology search of MMseqs2^41^ with AlphaFold2^42^. Predicted protein structures were searched against the Protein Data Bank database^43^ with Foldseek^44^. The top hits of PhnJ in both *Methylotenera* and *Methylomonadaceae* were experimentally validated PhnJ proteins in *Escherichia coli* K-12 (7Z19) ^45^ with e-values <10^-40^. 3D protein structure of the recovered PhnJ proteins was aligned with the validated PhnJ and visualized in ChimeraX v1.7.1. Positions of the four conserved cysteine residues and a glycine residue were highlighted.

### Identification of the methylphosphonate synthase MPnS

Reference sequences of the methylphosphonate synthase (MPnS) were obtained according as previously described^46^. All assembled contigs were searched against reference sequences with DIAMOND v2.0.9 using “-e 1e-5” ^47^. The extracted putative MPnS sequences were aligned with reference sequences using MAFFT v7.508 with default parameters^35^. Poorly aligned regions were removed with trimAl v1.4^36^. The phylogenetic tree was constructed with “-m MFP-B 1000” in IQ-TREE v2.2.0.3^37^ and visualized using iTOL v6.9 (https://itol.embl.de/). The presence of the 2-histidine-1-glutamine iron-coordinating triad and the two glutamine-adjacent residues (phenylalanine and isoleucine) required for methylphosphonate synthesis^46^ were manually checked based on the multiple sequence alignment (Supplementary dataset 1).

### Statistical analyses

The Pearson correlation coefficient (*R*) between methane concentration and relative abundance of *Methylotenera* was calculated in PASW Statistics v18.0. Genome size and coding density of *Methylotenera* in lake water and groundwater habitats were compared in t-tests in PASW Statistics v18.0. Statistical analyses were considered significant with *p*-values <0.05.

## Results and Discussion

### Biogeography of groundwater *Methylotenera*

Consistent with previous work on surface water^14–16^, methane concentration partially explained *Methylotenera* abundance (Figure 1a). What was striking about groundwater as a *Methylotenera* habitat, was that *Methylotenera* were so abundant, up to 88% relative sequence abundance in 16S rRNA gene amplicon libraries. To investigate the biogeography of *Methylotenera* in poorly connected groundwater habitats and discover what type of metabolisms contributed to their abundance, we analyzed shotgun metagenomes of twenty-six groundwater samples (Supplementary Table S1).

**Figure 1.**
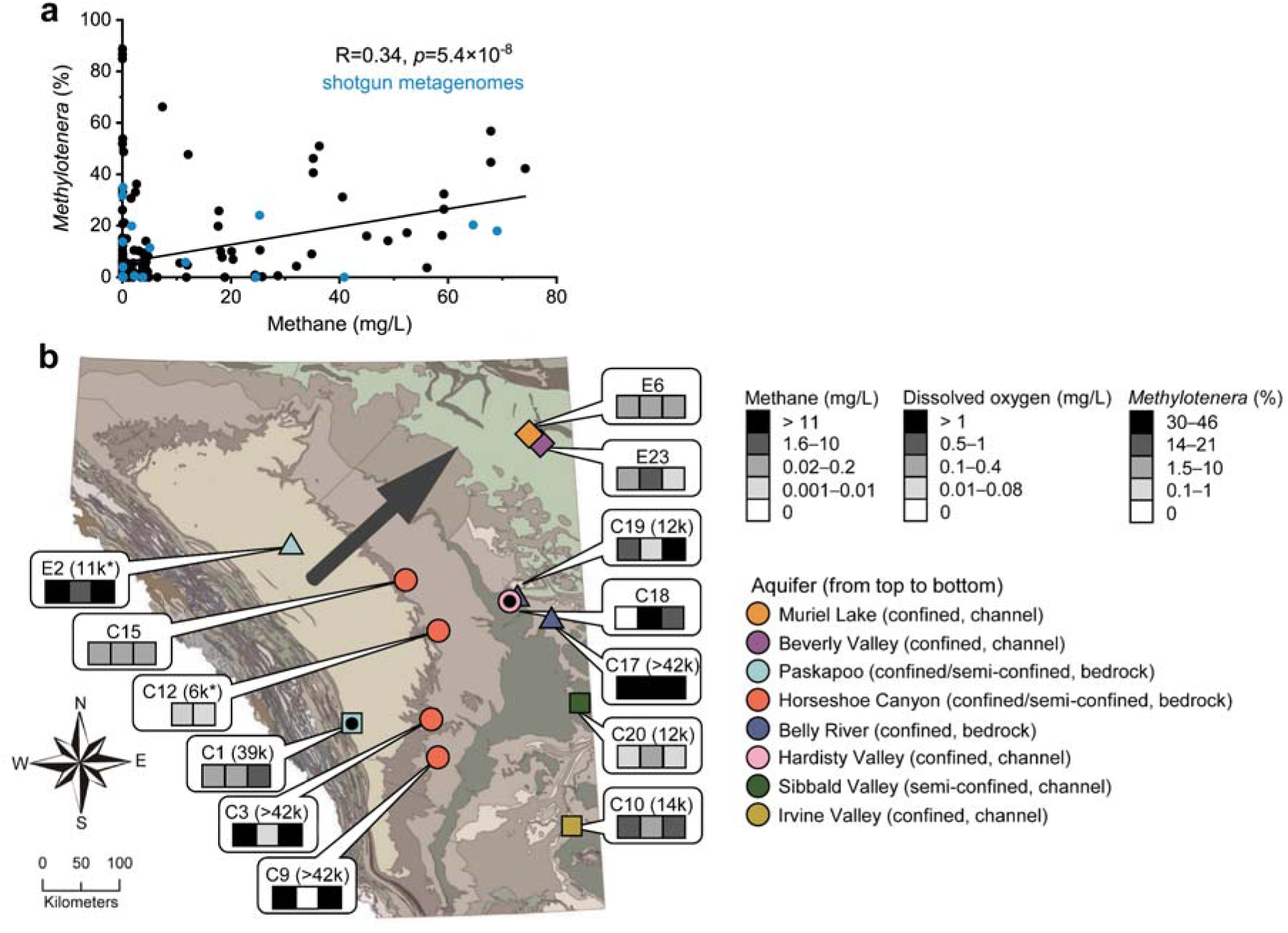
Wide dispersal of *Methylotenera* in unconnected groundwater habitats with highly variable methane and oxygen concentrations. (a) Methane concentrations partially explain *Methylotenera* abundances based on 16S rRNA gene amplicon data^17^. The shotgun-sequenced samples are in blue. (b) Occurrence of five distinct *Methylotenera* populations in Pleistocene-era groundwater with a range of oxygen and methane concentrations. Each population is indicated with a different shape: diamonds (99.994% average nucleotide identity, ANI), triangles (99.957%-99.985% ANI), circles (99.906%-99.999% ANI), squares (99.972%-99.996% ANI) and black dots (99.932% ANI). Numbers in brackets following sample IDs indicate groundwater age measured with ^14^C in dissolved inorganic carbon (DIC). * shows detection of ^3^H, an indicator for mixing with young water. The arrow indicates the general direction of groundwater flow, from the Rocky Mountains in the southwest towards the northeast. Marker colors indicate aquifer names. Aquifers are named after the hosting geological formation (background colors). The map was created with ArcGIS Pro v2.4.1. Thirteen out of twenty-six total samples (Supplementary Table S1) are shown on the map.

Thirty-six MAGs of *Methylotenera* were recovered (Supplementary Table S2). Twenty of these MAGs had estimated completeness >90% and contamination <5%. Based on sequencing reads mapping to these *Methylotenera* MAGs, the highest *Methylotenera* relative sequence abundance observed in the shotgun data was 35%, lower than observed with amplicon sequencing but still formidable compared to surface water habitats.

Among the recovered *Methylotenera* MAGs, there were five sets of nearly identical MAGs (>99.9% average nucleotide identity, Figure 1b, Supplementary Table S3). One might consider these five sets to represent five different *Methylotenera* populations. Based on estimates of bacterial mutation rates^31,32^, each of these populations emerged from a common ancestor between 1,000 and 94,000 years ago, for populations sharing 99.999% and 99.906% average nucleotide identity respectively. That is relatively recent in evolutionary terms. Sometimes MAGs from the same *Methylotenera* population were observed in the same aquifer. Surprisingly, they were also found in different aquifers that are currently not connected.

Based on radiocarbon and tritium dating (Supplementary Table S1), the groundwater in many of our samples recharged during the Pleistocene, during or before the last deglaciation (12,000 – 115,000 years ago) ^48^, with occasional contributions of much younger water (Figure 1b). Groundwater recharge was much more active during ice ages and the >2 km thick Laurentide ice sheet that covered much of Canada east of our study area may have altered or reversed groundwater flow directions^49^. During the Pleistocene, aquifers that are currently isolated from each other may have been connected. Supercharged groundwater flow could explain colonization of presently unconnected aquifers by the observed *Methylotenera* populations. Some of these populations must have lived isolated from each other and the surface for tens of thousands of years.

When the same aquifer was sampled at multiple locations, very different concentrations of oxygen and methane were measured, indicating that each aquifer contains different environments, likely impacted by local bedrock features such as deposits of shale and pyrite. Thus, each aquifer may offer multiple ecological niches that could favor different *Methylotenera* populations. Methane and oxygen were often present together, just like in the upper oceans. In the groundwater though, methane concentrations were always higher. Based on the isotopic signature, methane mainly originated from methanogenesis^18^. The presence of “dark oxygen” in ancient waters has been noted before^18^. It may indicate active oxygen production as it is hard to explain how oxygen could persist for thousands of years in the presence of methane; in addition to *Methylotenera*, these aquifers also hosted methane-oxidizing *Methylomonadaceae* (data not shown).

### Phylogeny of groundwater *Methylotenera*

*Methylotenera* phylogeny was analyzed based on 71 bacterial marker genes and 88 reference genomes publicly available in GTDB. The resulting phylogenetic tree featured five clades associated with specific habitats (Figure 2a, Supplementary Table S4). Two of those (LW1, LW2) were made up exclusively of genomes found in the water column of lakes. Thirty-six of the groundwater MAGs formed three clades, G1, G2 and G3, which also contained some genomes previously obtained from groundwater, groundwater-associated habitats or sediments.

**Figure 2.**
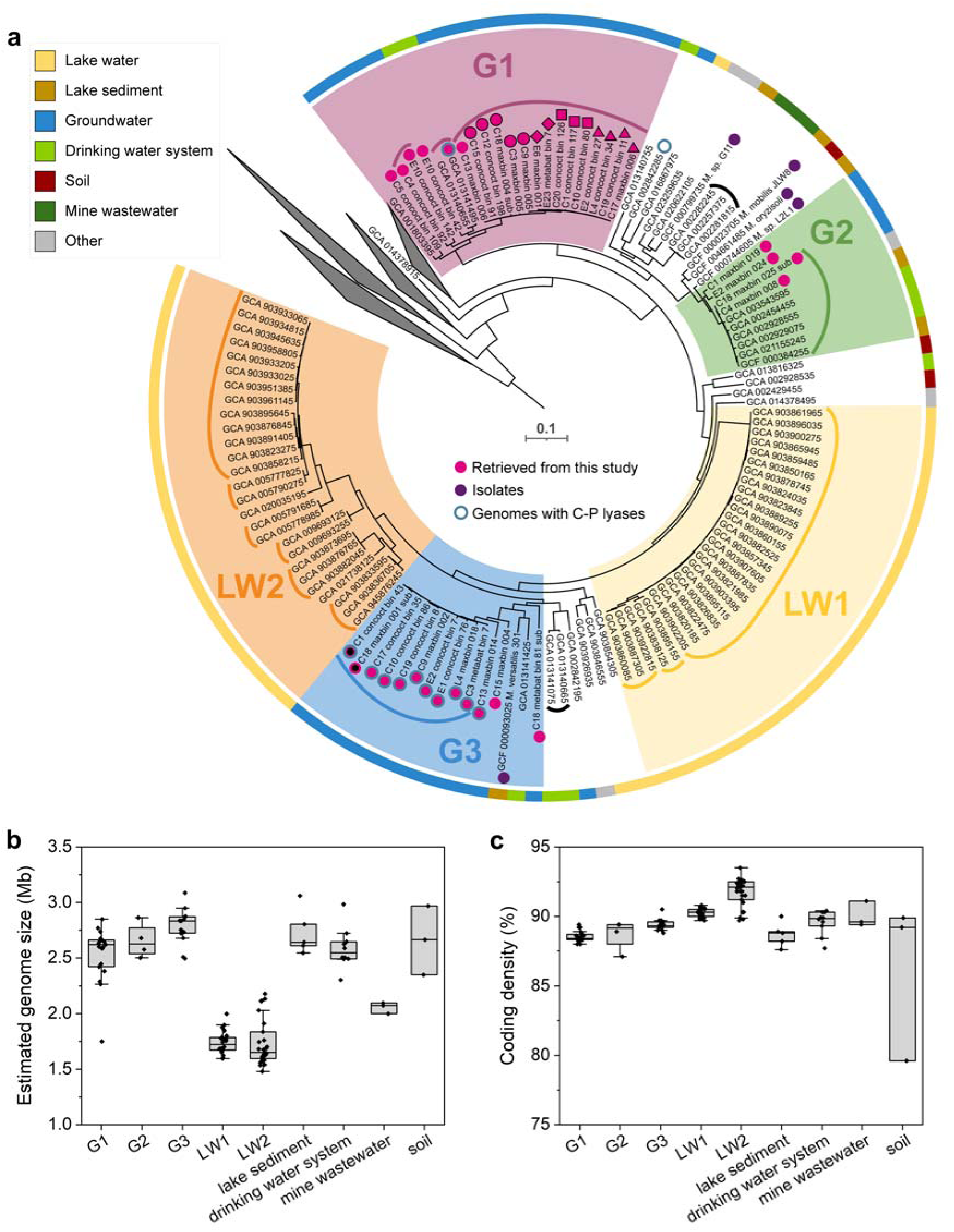
Groundwater and lake water *Methylotenera* have distinct phylogeny and genome size. (a) Phylogenetic tree of *Methylotenera* MAGs recovered in this study (pink markers) and reference genomes from GTDB including those of isolated strains (purple circles). Curves show groups of genomes that share 95+% average nucleotide identity (species). Black thick edges around the circles indicate the presence of C-P lyase genes involved in methylphosphonate breakdown (see Figure 4). Marker shapes correspond to the five populations in Figure 1b. The ring around the tree indicates habitat. (b) Estimated genome size and (c) coding density of *Methylotenera* genomes recovered from different environments. Both MAG and sample (Figure 1b) identifiers are indicated for each recovered genome.

G1 consisted of nineteen groundwater MAGs and three reference genomes, one from a groundwater well in Colorado, USA, and the other two recovered from a sand filter for a drinking water treatment plant in the Netherlands. G1 genomes shared >86% average nucleotide identity (ANI) and comprised seven different species-level clades, each sharing >97% ANI^50^, and four of the populations shown in Figure 1b.

G2 consisted of four groundwater MAGs as well as six reference genomes, two from lake sediments, two from a groundwater-derived drinking water system, one from a hydrocarbon enrichment culture and one from a thick biofilm clogging water pumps in an ice machine. G2 genomes shared >87% ANI and comprised three different species-level clades. *Methylotenera oryzisoli* (GCF_004661485), isolated from the rice rhizosphere^7^, *Methylotenera mobilis* JLW8 (GCF_000023705) ^5^ and *Methylotenera* sp. L2L1 (GCF_000744605) ^1^, both isolated from Lake Washington sediments, formed a sister clade to G2. *Methylotenera* sp. G11 (GCF_ 000799735) ^1^, also from Lake Washington sediments, was at the root of G2. As we will show below, in groundwater G2 is less abundant than both G1 and G3 and mainly found in water with low methane concentrations (see below).

G3 consisted of thirteen groundwater MAGs and two reference genomes, one from a lake sediment and one from the above-mentioned sand-filter. G3 genomes shared >78% ANI and comprised six different species-level clades and one population in Figure 1b. *Methylotenera versatilis* 301 (GCF_000093025) ^6^, isolated from Lake Washington sediments, was part of G3. As we will show below, G3 *Methylotenera* are associated with high methane concentrations and aerobic methane production from methylphosphonate.

Genomes recovered from the water columns of lakes were mainly within LW1 and LW2, with only three exceptions (Figure 2a). They were mostly retrieved from a comprehensive dataset of metagenomes from stratified freshwater lakes in Sweden, Finland, Switzerland and other countries^51^. LW1 and LW2 had significantly smaller genome size and higher coding density than groundwater *Methylotenera* clades (*p*<0.05, Figure 2b). This indicated that surface water *Methylotenera* were experiencing genome streamlining^52^, which was previously observed within the same family (*Methylophilaceae*) for two pelagic lineages, Ca. *Methylopumilus* and OM43^29^. Our analysis (Figure 2a) showed that the ancestors of G3 *Methylotenera* colonized surface waters at least twice, each time resulting in massive gene loss.

Apparently, surface water environments select for *Methylotenera* with limited, specialized metabolisms living on minimized energy and/or nutrient budgets. In contrast, groundwater *Methylotenera* are able to maintain a much larger accessory genome, leaving more room for metabolic flexibility and other cellular features. The intertwined evolutionary histories of groundwater and sediment *Methylotenera* points at overlaps in ecological niche space between these two structured habitats.

### Metabolisms of groundwater and lake water *Methylotenera*

To explore if G1-3 might occupy distinct ecological niches and how those niches were different from LW1-2 lake *Methylotenera*, we compared gene content of twenty-seven high-quality MAGs associated with G1-3, LW1-2. Expression of genes by groundwater *Methylotenera* was inferred from proteomics analysis of five groundwater samples (Figure 3, Supplementary Table S5, S6).

**Figure 3.**
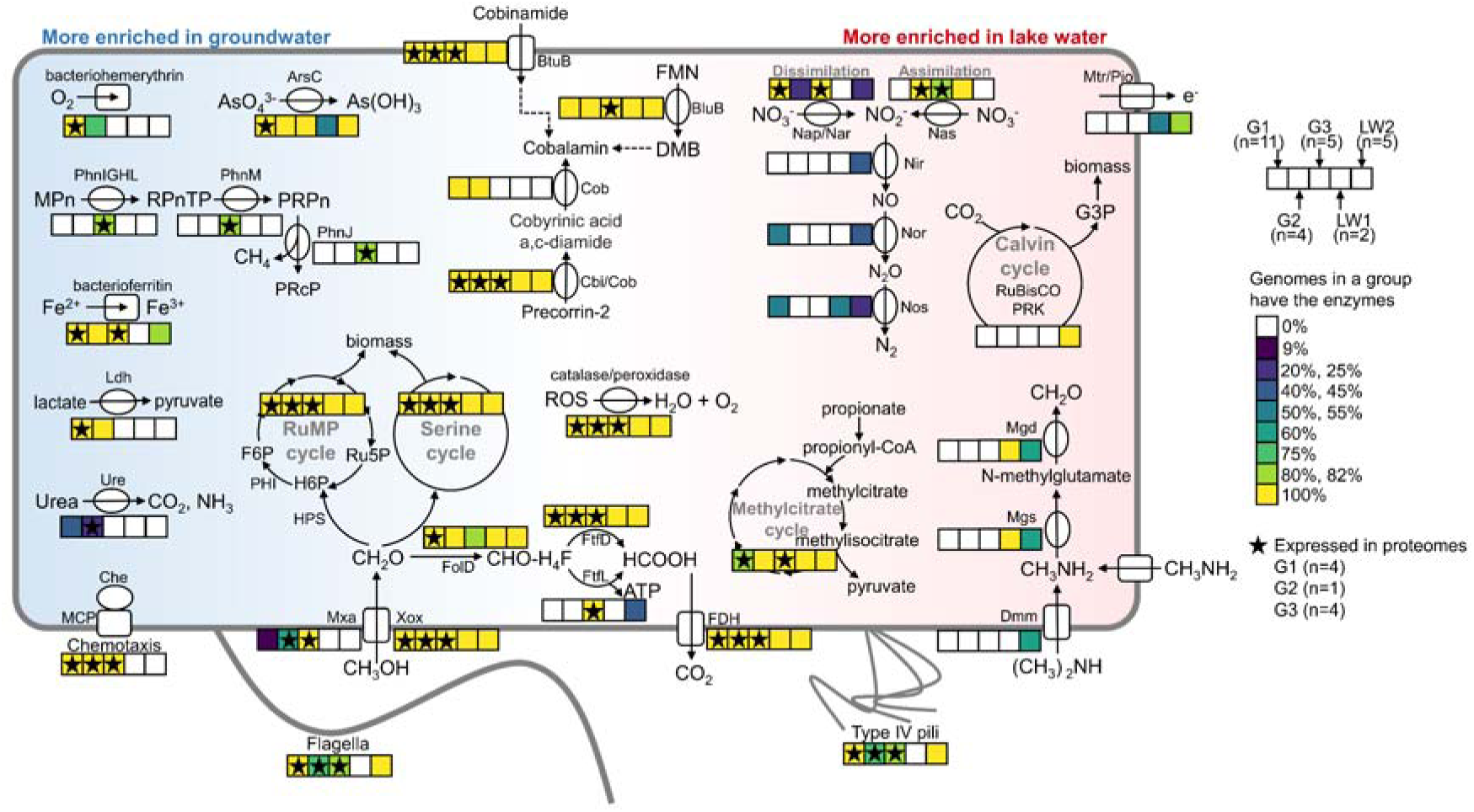
Metabolisms of *Methylotenera* in groundwater and lake water habitats. The heatmaps indicate the presence of associated genes in high-quality genomes of G1 (n=11), G2 (n=4), G3 (n=5), LW1 (n=2) and LW2 (n=5). Pathways with at least one gene expressed in proteomes are indicated with stars. The metabolic pathways more enriched in *Methylotenera* genomes sampled from groundwater are shown on the left. Genes enriched in lake water are shown on the right. Abbreviations as follows: methylphosphonate (MPn), alpha-D-ribose 1-methylphosphonate 5-triphosphate (RPnTP), alpha-D-ribose-1-phosphonate-5-phosphate (PRPn), alpha-D-ribose-1,2-cyclic-phosphate-5-phosphate (PRcP), flavin mononucleotide (FMN), 5,6-dimethylbenzimidazole (DMB), RuMP cycle (ribulose monophosphate cycle), hexulose-6-phosphate (H6P), fructose 6-phosphate (F6P), ribulose 5-phosphate (Ru5P), formyl-tetrahydrofolate (CHO-H_4_F), glyceraldehyde-3-phosphate (G3P), reactive oxygen species (ROS). The more detailed information is presented in Supplementary Table S5.

Key enzymes involved in the Calvin cycle, RubisCO and phosphoribulokinase, were exclusively observed in all LW2 genomes, indicating that only those lake water *Methylotenera* were facultative autotrophs. On the other hand, lactate dehydrogenase was exclusively observed in all G1-2 genomes, indicating the potential for fermentative lactic acid production by groundwater *Methylotenera*. Urease was enriched in some G1-G2 genomes but absent in LW1-2, indicating that urea was a relevant nitrogen source for groundwater but not lake water *Methylotenera*. The decaheme-associated outer membrane protein MtrB/PioB for extracellular electron transport was exclusively present in some LW1-2 genomes.

LW1 did not have genes for flagella and type IV pili, whereas all other clades had. Genes involved in chemotaxis were present and expressed in all G1-3 genomes but not in LW1-2. As both groundwater and sediment habitats are more structured than lake water, motility and chemotaxis could be essential to cope with spatial gradients of oxygen and nutrients.

Bacteriohemerythrin was observed in all G1 genomes and three G2 genomes. Expression of bacteriohemerythrin was also observed. It was not present in LW1-2. Bacteriohemerythrin is used to bind oxygen, for storage, enzyme delivery and sensing^53^. These functions could be important in a habitat with spatial oxygen gradients or temporal oxygen fluctuations. In lake water, oxygen levels might be more stable. Defense against reactive oxygen species (ROS) was still important in groundwater habitats, as catalase, peroxiredoxin, thioredoxin, superoxide dismutase and peroxidase were present in all G1-3 genomes. ROS defense proteins were actively expressed in groundwater-the total relative abundance of ROS defense proteins was about 1% to 4% in *Methylotenera* proteomes.

As for cobalamin synthesis, most core genes were present in all G1-2 genomes. All G3, LW1-2 contained genes for corrin ring biosynthesis but lacked those involved in following steps, final synthesis and repair of cobalamin^30^. All genomes had the cobalamin transporter, BtuB, a TonB-dependent outer membrane cobalamin receptor and transporter, which was expressed as well, indicating these populations could also acquire it from their environment. All genomes contained 5,6-dimethylbenzimidazole (DMB) synthase, BluB, which is involved in synthesis and activation of DMB, the cobalamin lower ligand. The expression of BluB was only detected in G3, indicating G3 synthesized cobalamin from DMB.

With regard to methylotrophy, the lanthanide-dependent methanol dehydrogenase Xox was present in all genomes, while the calcium-dependent methanol dehydrogenase was only present among some G1-2 and all G3 genomes. All genomes lacked enzymes for direct conversion of formaldehyde to formate. Instead they used the tetrahydrofolate (THF)-dependent pathway^54^. In this pathway, formaldehyde first binds to tetrahydrofolate (H_4_F) spontaneously to form methylene-tetrahydrofolate (CH_2_-H_4_F), then the enzyme FolD converts CH_2_-H_4_F to methenyl-tetrahydrofolate (CH-H_4_F), which is subsequently converted to formyl-tetrahydrofolate (CHO-H_4_F). This product is then converted to H_4_F and formate by formyltetrahydrofolate deformylase, which was present in all genomes, or with formate-tetrahydrofolate ligase, which was only present in all G3 and some LW2 genomes. Formate-tetrahydrofolate ligase produces ATP, while formyltetrahydrofolate deformylase does not, so G3 and LW2 might conserve additional energy during formaldehyde oxidation. Formate dehydrogenase, the ribulose monophosphate pathway and the serine pathway for formaldehyde assimilation were present in all genomes. The methylcitrate cycle for oxidizing propionate to pyruvate via methylcitrate was present in most genomes^55^. Apart from methanol, formaldehyde and formate, LW1 and LW2 were able to use methylamine, while three LW2 populations could also use dimethylamine.

Therefore, groundwater *Methylotenera* were able to use methanol, formaldehyde and formate, which were most likely provided by the accompanying methanotroph *Methylomonadaceae*. The co-existence of methanotrophic *Methylobacter* (*Methylomonadaceae*) and methylotrophic *Methylotenera* was previously observed in a methane-rich shallow aquifer^56^, lake water^57^ and sediments^12,13^. The cross feeding was suggested to be regulated by the expression of the lanthanide-dependent Xox-versus the calcium-dependent Mxa-type methanol dehydrogenase in *Methylobacter*: the expression of Mxa with lower affinity for methanol leads to a loss of methanol to the external surroundings of cells which benefits *Methylotenera*. Excretion of a lanthanide-scavenging molecule by *Methylotenera* was shown to play a role in this interaction^58^. In the groundwater, we found both Mxa and Xox were expressed by *Methylomonadaceae* (Supplementary Table S5), indicating that they do have access to lanthanides.

Alternatively, denitrifying *Methylotenera* could mop up C1 compounds that could not be oxidized further by oxygen-limited *Methylomonadaceae*^16^. Although previous studies showed the importance of nitrate as the electron acceptor in driving methylotrophic metabolism in *Methylotenera*^15,16^, we found denitrification was not so prevalent in *Methylotenera*, neither in lake water nor groundwater. While the dissimilatory nitrate reductase Nar was present in most genomes, enzymes in subsequent steps of denitrification were less prevalent: the nitrite reductase NirK was only present in two LW2 genomes, co-existing with the nitric-oxide reductase Nor, while one of them also has the nitrous-oxide reductase NosZ. Nor and NosZ co-existed in six G1 genomes, and NosZ was also observed in a LW1 genome. Among all denitrifying enzymes, only Nar was expressed in proteomes, which might be the driver for methanol cross-feeding. Fermentation and arsenate reduction could also drive cross-feeding in groundwater *Methylotenera*.

### Aerobic methane production from methylphosphonate

Interestingly, we detected a pathway for methylphosphonate assimilation in G3 (Figure 3). In this pathway, the C-P lyase core complex Phn converts methylphosphonate into methane and phosphate as follows: First PhnIGHL converts methylphosphonate and ATP into alpha-D-ribose 1-methylphosphonate 5-triphosphate (RPnTP) and adenine. Next, PhnM cleaves off pyrophosphate resulting in alpha-D-ribose-1-phosphonate-5-phosphate (PRPn). Finally, PhnJ generates alpha-D-ribose-1,2-cyclic-phosphate-5-phosphate (PRcP) releasing methane. The pathway was present in nine genomes that formed a new species in G3. This species also featured one MAG (C18_maxbin.001_sub, Figure 1a) that did not encode the C-P lyase core complex itself, but it did feature a short contig with an incomplete phosphonate metabolism transcriptional regulator PhnF next to PtsS, ArsBC, RF-2 and LysRS, the same gene arrangement found adjacent to the C-P lyase core complex genes in the other G3 *Methylotenera* genomes (Figure 4a). Thus, the C-P lyase appears to be a defining feature of this G3 species, and proteomics showed that it was actively expressed (Figure 4, 5, Supplementary Table S6).

**Figure 4.**
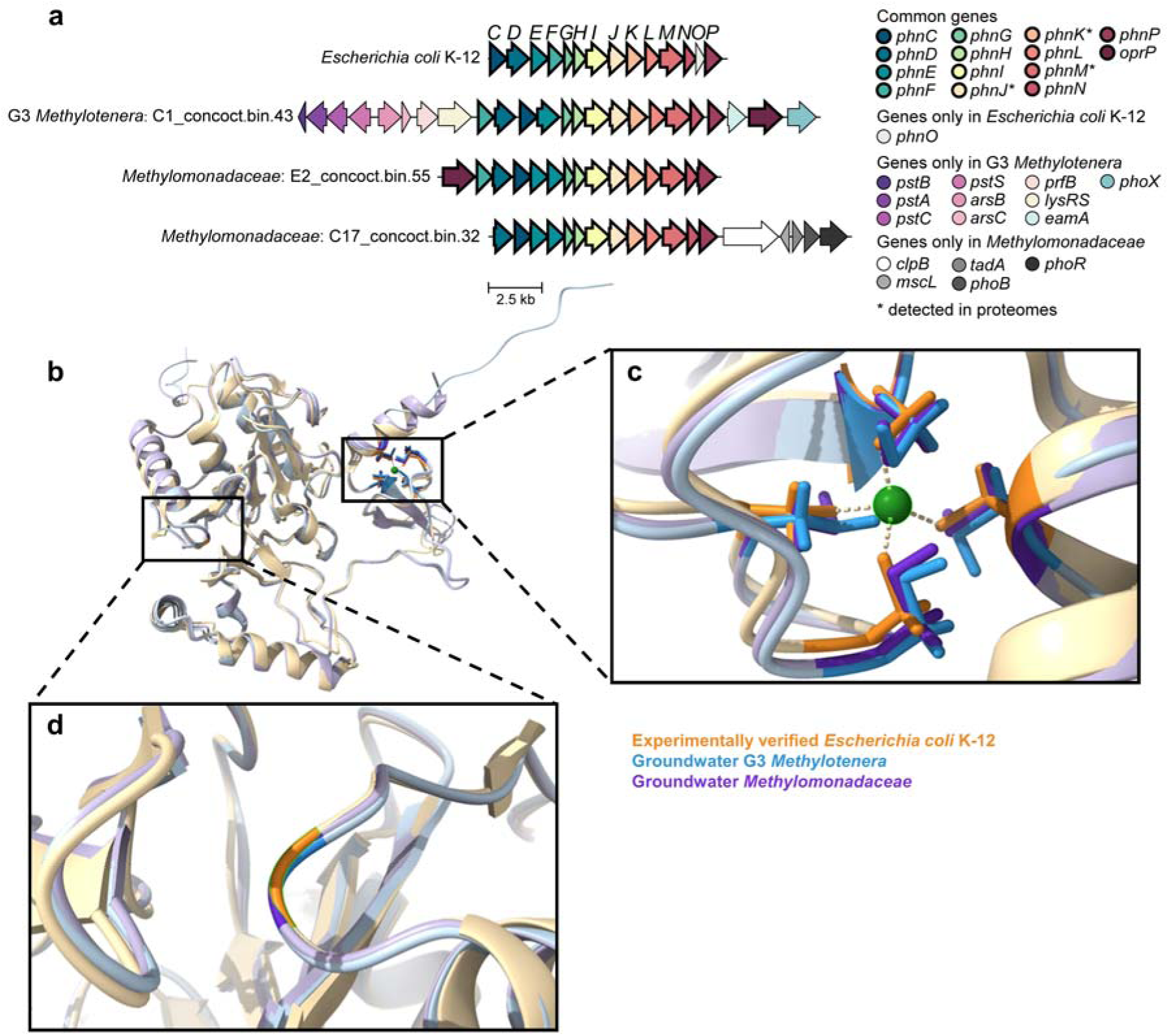
Conservation of essential genes, protein structure and amino acids needed for methane production by G3 *Methylotenera* and *Methylomonadaceae*. (a) Gene arrangement of the Phn operon and flanking genes in experimentally verified *Escherichia coli* K-12, G3 *Methylotenera* and *Methylomonadaceae* MAGs. The letters at the top indicate subunits of the Phn complex. * indicates gene expression was detected in proteomes. (b) Protein structure models of PhnJ in G3 *Methylotenera* (blue) and *Methylomonadaceae* (purple) aligned with *E. coli* PhnJ (protein data bank 7Z19, gold). Zoomed views of the conserved four cysteine residues and glycine-32 are shown in (c) and (d). Conserved residues are shown in corresponding dark colors. The zinc ion is shown in green. Gene abbreviations in (a) as follows: phosphonate ABC transporter ATP-binding protein (*phnC*), phosphonate ABC transporter substrate-binding protein (*phnD*), phosphonate ABC transporter permease protein (*phnE*), phosphonate metabolism transcriptional regulator (*phnF*), phosphonate C-P lyase system protein subunit GHIJKLM (*phnGHIJKLM*), phosphonate metabolism protein/1,5-bisphosphokinase (PRPP-forming) (*phnN*), aminophosphonate N-acetyl-transferase (*phnO*), carbon-phosphorus lyase complex accessory protein (*phnP*), phosphate-selective porin (*oprP*), phosphate ABC transporter ATP-binding protein (*pstB*), phosphate ABC transporter permease protein (*pstA*), phosphate ABC transporter permease protein (*pstC*), phosphate ABC transporter substrate-binding protein (*pstS*), arsenite efflux pump (*arsB*), arsenate reductase (*arsC*), peptide chain release factor 2 (*prfB*), lysine--tRNA ligase (*lysRS*), phosphonate utilization associated putative membrane protein (*eamA*), alkaline phosphatase (*phoX*), ATP-dependent chaperone (*clpB*), large-conductance mechanosensitive channel protein (*mscL*), tRNA(Arg) A34 adenosine deaminase (*tadA*), phosphate regulon transcriptional regulator (*phoB*), phosphate regulon sensor histidine kinase (*phoR*).

We explored if, among the bacteria in the investigated groundwater samples, methylphosphonate metabolism was unique to G3 *Methylotenera*. It appeared that *Methylotenera*’s methane-oxidizing partners, *Methylomonadaceae*, also encoded and expressed PhnJ, as well as chemolithoautotrophic *Hydrogenophaga* and many other bacteria often found in our groundwater samples, such as *Phenylobacterium*, *Polaromonas* and *Rhodoferax* (Supplementary Table S7).

Figure 4a compares the Phn gene clusters found in G3 *Methylotenera* and *Methylomonadaceae* to the experimentally validated *Escherichia coli* gene cluster. All genes were observed in G3 *Methylotenera* and *Methylomonadaceae*, except for PhnO which is only required in the catabolism of 1-aminoalkylphosphonic acid^59^. PhnGHIJKLMNP in *Methylotenera* and *Methylomonadaceae* were in the same order as those in *E. coli*. For PhnCDEF in *E*. *coli*, the order of these genes was PhnFDCE in G3 *Methylotenera.* In *Methylomonadaceaea*, sometimes PhnF was not present, while PhnD was ahead of PhnC followed by two PhnE. The 3D structure of the key enzyme for methane production from methylphosphonate, PhnJ, was modeled and inspected (Figure 4b). The presence of the four conserved cysteine residues (Cys241, 244, 266 and 272) in PhnJ of both G3 *Methylotenera* and *Methylomonadaceae* were confirmed (Figure 4c). These participate in the binding of an iron-sulfur cluster and the coordination of the zinc atom. During the catalytic cycle of the enzyme, after activation by S-adenosyl-methionine, Cys272 and Gly32 (also conserved, Figure 4d) donate and regenerate the H· radical needed for methane production^60^. We also compared the phylogeny of PhnJ to the *E. coli* protein (Figure S1). The G3 *Methylotenera* and *Methylomonadaceae* proteins were closely related, sharing 80-82% amino acid identity. They were both in one of two major clades within this protein family, whereas the validated *E. coli* protein was in the other clade. In the ocean, PhnJ proteins from both clades were observed, while the clade with *E. coli* was more abundant and most PhnJ proteins were found in members of *Alphaproteobacteria*, such as SAR116 and *Pelagibacter*^61–63^.

Breaking the stable carbon-phosphorus bond by the Phn complex requires four ATP^45^, with an additional ATP consumed in binding the methylphosphonate itself. Oxidation of the produced methane would not yield sufficient energy to compensate for these energy expenses. However, the energy expenses are justified when growth is phosphate-limited^21,61^ and the aim is to assimilate phosphonate. The genes of the Phn complex were located next to genes encoding the phosphate ABC transporter in G3 *Methylotenera* and next to the Pho phosphate regulon in *Methylomonadaceae* (Figure 4a). These genes are involved in sensing and transport of phosphate^64^. An alkaline phosphatase gene, *phoX*, was also situated close to the Phn genes in G3 *Methylotenera*. PhoX is used to assimilate phosphate from organic compounds^65^. Together, these findings strongly suggested that PhnJ was expressed in response to phosphate limitation. We did not detect a relationship between the relative abundance of microorganisms with PhnJ and phosphate concentration (Figure 5b, Supplementary Table S1). However, as mentioned before, subsurface habitats are structured and productive biofilms can be phosphate-limited even though the groundwater might still contain some phosphate^18^.

**Figure 5.**
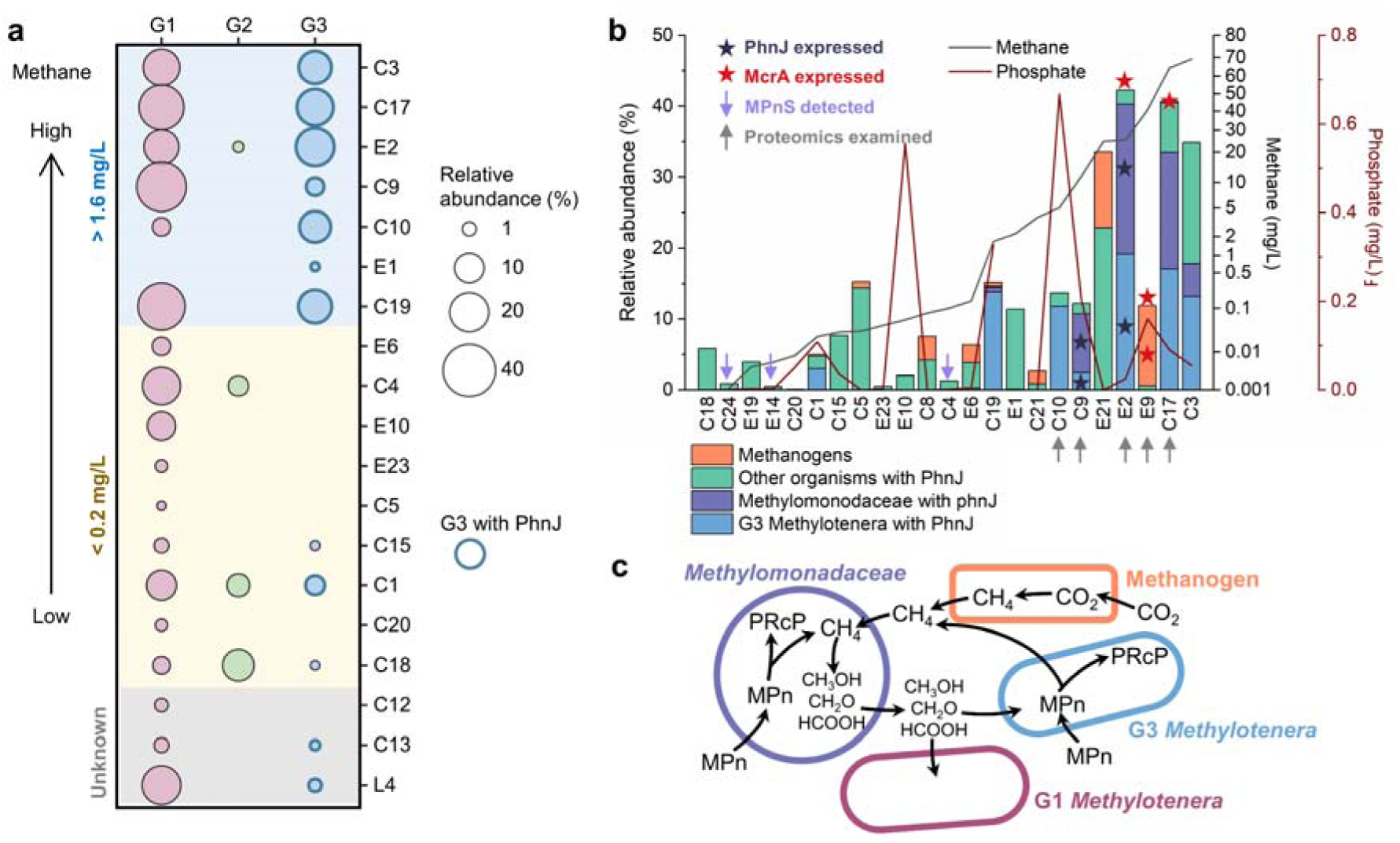
Methane-rich groundwater is associated with methane-producing *Methylotenera*. (a) Relative abundance of G3 (*R*=0.598, *p*=0.003) but not G1 (*R*=0.383, *p*=0.072) and G2 *Methylotenera* (*R*=-0.176, *p*=0.421) in metagenomes correlates with methane concentration (Supplementary Table S1, 2). G3 *Methylotenera* with PhnJ are indicated with thick edges around the circles. (b) Relative abundance of both methanogens and microorganisms encoding the C-P lyase core complex increased with methane concentrations, while there was no correlation with phosphate concentrations (phosphate concentration in E1 is unknown). Arrows below sample identifiers indicate samples examined with proteomics. Purple arrows indicate the detection of methylphosphonate synthase, MPnS. Dark blue stars indicate the expression of PhnJ. Red stars indicate the expression of methyl-coenzyme M reductase subunit alpha McrA. (c) Proposed metabolic handshakes between methanogens, methanotrophic *Methylomonodaceae* and methylotrophic *Methylotenera*.

Methylphosphonate is an intermediate in the breakdown of natural and man-made organic phosphonates. Studies of the marine phosphorus cycle have estimated that phosphonates make up about 10-25% of the high molecular weight dissolved organic phosphorus compounds^66,67^. These phosphonates can decorate lipids, polysaccharides and proteins. Methylphosphonate is produced with the methylphosphonate synthase MPnS in microorganisms^46,68^. MPnS was detected in only three samples, C4, C24 and E14 (Figure 5b, Figure S2). Methylphosphonate might also be derived from the breakdown of glyphosate, an important herbicide in agriculture and forestry^64^. However, as many of the investigated samples were from confined aquifers with water that infiltrated before the start of the Holocene, we would not expect herbicides to be relevant. Future research could address if groundwater harbors a geological source of methylphosphonate, and if so, what would be its isotopic composition. Another fascinating possibility is the presence of a natural phosphonate-based antimicrobial in groundwater that prompts expression of defensive degradation.

Among groundwater *Methylotenera*, high abundance G3 with PhnJ was strongly associated with methane-rich groundwater (*R*=0.598, *p*=0.003, Figure 5a, Figure 1). G1 was enriched in water with high methane concentrations but was also present at lower methane concentrations. In contrast, G2 was mainly found in water with low methane concentrations. To explore if methylphosphonate-degrading *Methylotenera* were also associated with high methane concentrations in other studies, we checked the presence of the Phn complex in all *Methylotenera* genomes of Figure 1a. Phn was also found in two other *Methylotenera* MAGs: GCA_002842285 (in the sister group of G2) and GCA_013141495 (in G1). GCA_002842285 was recovered from a groundwater well with methane making up 99.4% of the headspace gas^69^. GCA_013141495 was from a primary rapid sand filter fed with groundwater rich in methane (5.2 mg/L) ^70^. Similar to G3, the latter co-existed with several *Methylomonadaceae* populations^70^.

To explore if methylphosphonate could potentially be a significant source of methane, we compared the relative abundances of methanogens and of PhnJ-containing microbes in methane-rich samples (Figure 5b). Methanogens were somewhat abundant in some of the samples with high methane concentrations (e.g. E9, E21). In other samples, methanogens were barely detectable and the relative abundance of methylphosphonate degraders was much higher (e.g. C3, C17, E2, C9).

The five proteomes showed active expression of both McrA in methanogens, and PhnJ in two G3 *Methylotenera* and four *Methylomonadaceae* populations. Although G3 *Methylotenera* are unable to use methane, the produced methane might be oxidized by aerobic methanotroph *Methylomonadaceae*. *Methylotenera* could still benefit from methylphosphonate mineralization by cross-feeding off intermediates of methane oxidation, such as methanol, formaldehyde and formate (Figure 5c). As mentioned before, the large amount of energy needed to split the carbon-phosphorous bond could never be recovered by oxidizing the produced methane, so this strategy would not be able to drive net growth. Further, although methylphosphonate degradation could contribute to methane production, the isotopic signature of the methane in these samples was still consistent with methanogenesis as the main pathway of methane production^18^. Therefore, it is more likely that the correlation between methane concentration and Phn abundance can be explained by high microbial productivity in these waters^18^, resulting in phosphate limitation.

## Data availability

Amplicons in this study are under the Bioproject PRJNA861683 and PRJNA700657. Metagenomes and metagenome-assembled-genomes are under the Bioproject PRJNA700657 (NCBI). The mass spectrometry proteomics data have been deposited to the ProteomeXchange Consortium via the PRIDE partner repository ^71^ with the dataset identifier PXD044305.

## Acknowledgements

The authors thank the Groundwater Observation Well Network team members of Alberta Environment and Protected Areas (https://www.alberta.ca/groundwater-observation-well-network.aspx) for providing access to groundwater monitoring wells, for sampling and providing highest quality groundwater samples, and for sharing measurement results and expertise. Financial support from Alberta Innovates through its Water Innovation Program is gratefully acknowledged. We would like to thank the University of Calgary’s Center for Health Genomics and Informatics for sequencing and informatics services. This study was supported by the Natural Sciences and Engineering Research Council (NSERC) through Discovery Grants to Marc Strous and Bernhard Mayer, Canada Research Chair (CRC-2020-00257, MS) to Marc Strous, the Canada Foundation for Innovation (CFI), the Digital Research Alliance of Canada, the Canada First Research Excellence Fund (CFREF), the Government of Alberta, and the University of Calgary.

## Author contributions

SL, MS and MD designed the study. SL, XD, PH, JB, JB, CM, BM, MS and MD analyzed the data. SL, MS and MD wrote the manuscript.

