## Supplementary Information for "Aerobic methane production by methylotrophic *Methylotenera* in groundwater"

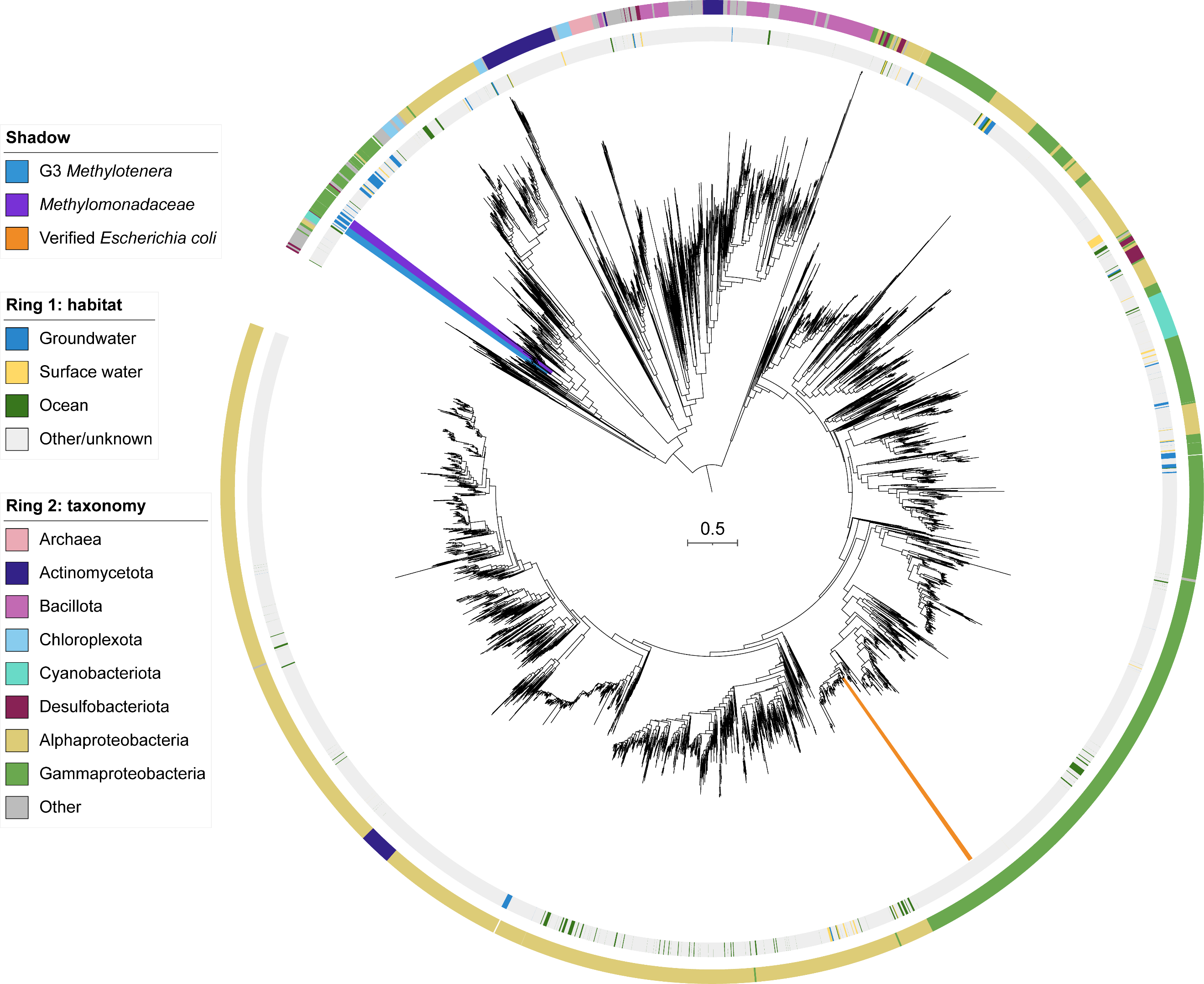

Figure S1. **Phylogenetic tree of the C-P lyase core complex subunit J (PhnJ)**. PhnJ sequences retrieved from this study and the reference database are included (Supplementary Table S7). Blue and purple shadows on leaves indicate those from G3 *Methylotenera* and *Methylomonadaceae* which are mostly retrieved from this study. Gold shadow indicates those experimentally verified PhnJ sequences in *Escherichia* *coli*. From inside to outside, the two rings around the tree indicate: (1) habitat and (2) taxonomy of the genome.

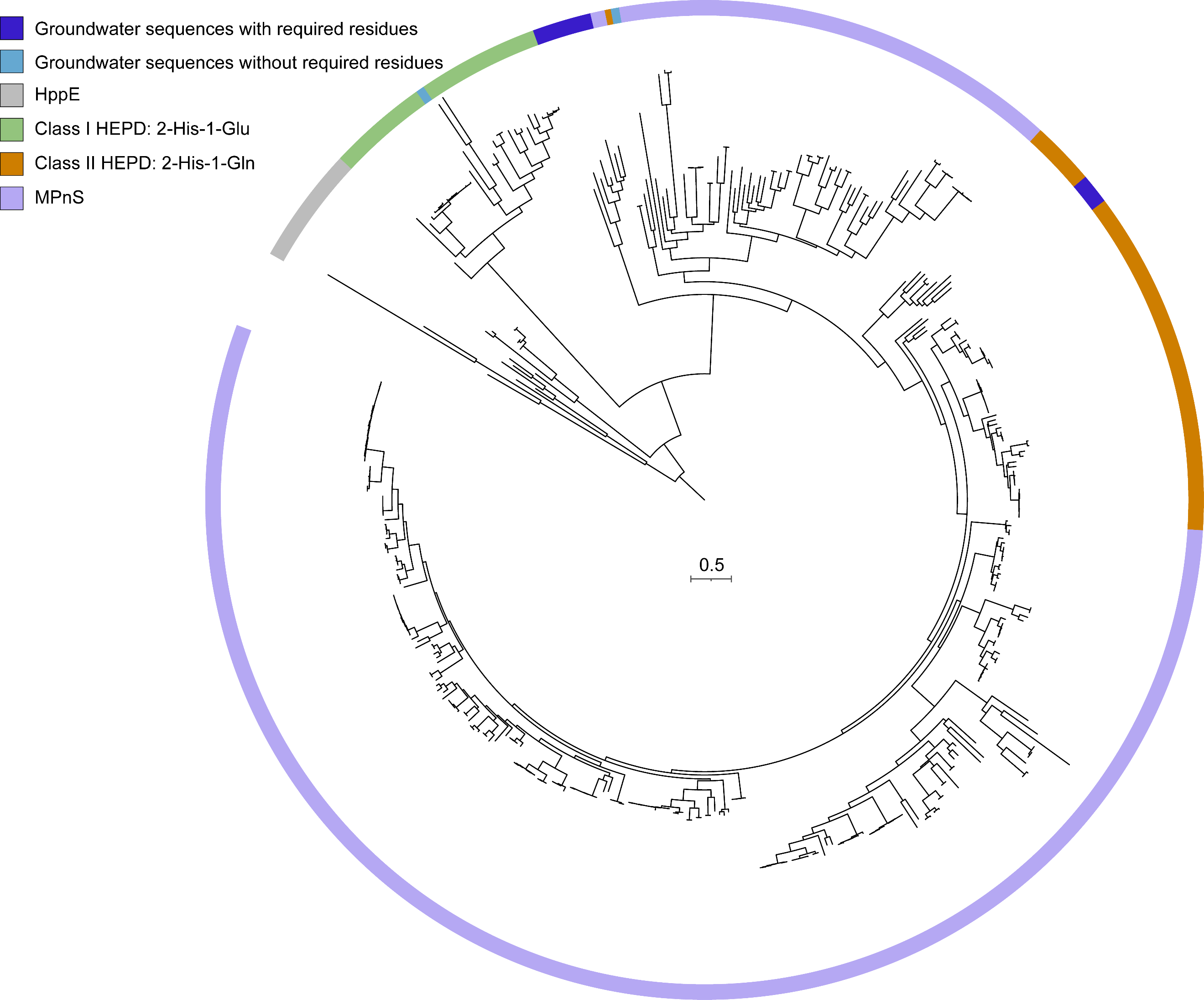

Figure S2. **Phylogenetic tree of the methylphosphonate synthase (MPnS)**. MPnS like sequences retrieved from this study and the reference sequences obtained from a previous study (Born et al., 2017) are included (Supplementary Table S8). The presence of the 2-histidine-1-glutamine iron-coordinating triad and the two glutamine-adjacent residues (phenylalanine and isoleucine) required for methylphosphonate synthesis (Born et al., 2017) were checked (Supplementary dataset 1). The ring around the tree indicates the groundwater sequences and the annotated reference sequences.
